# A Tale of Two Parasites: A Glimpse into the RNA Methylome of Patient-derived *Plasmodium falciparum* and *Plasmodium vivax* isolates

**DOI:** 10.1101/2023.12.26.573234

**Authors:** Priyanka Roy, Sukriti Gujarati, Pallavi Gupta, Ishaan Gupta, Tanmaya Mahapatra, Dinesh Gupta, Sanjay Kumar Kochar, Dhanpat Kumar Kochar, Ashis Das

## Abstract

**Background:** Understanding the molecular mechanisms of the malarial parasites in hosts is crucial for developing effective treatments. Epitranscriptomic research on pathogens has unveiled the significance of RNA methylation in gene regulation and pathogenesis. We present the first report investigating methylation signatures and alternative splicing events using Nanopore Direct RNA Sequencing to single-base resolution in *Plasmodium falciparum* and *P. vivax* clinical isolates with hepatic dysfunction complications.

**Methodology:** We performed direct RNA Sequencing using Nanopore from clinical isolates of *P. falciparum* and *P. vivax* showing hepatic dysfunction manifestation. Subsequently we performed transcriptome reconstruction using FLAIR and transcript classification using SQANTI3 followed by methylation detection using CHEUI and m6Anet to identify N6-methyladenosine (m6A) and 5-methylcytosine (m5C) methylation signatures. We also documented alternative splicing events from both the datasets.

**Results:** The reference genome of *Plasmodium* reports >5000 genes out of which we have identified ∼50% as expressed in the two sequenced isolates, including novel isoforms and intergenic transcripts, highlighting extensive transcriptome diversity. The distinct RNA methylation profiles of m^6^A and m^5^C from the expressed transcripts were observed in sense, Natural Antisense Transcripts (NATs) and intergenic categories hinting at species-specific regulatory mechanisms. Modified transcripts originating from apicoplast and mitochondrial genomes have also been detected. These modifications are unevenly present in the annotated regions of the mRNA, potentially influencing mRNA export and translation. We have observed several splicing events, with alternative 3’and 5’end splicing predominating in our datasets suggesting differences in translational kinetics and possible protein characteristics in these disease conditions.

**Conclusion:** In our data we are showing the presence of modified sense, NATs and alternatively spliced transcripts. These phenomena together suggest the presence of multiple regulatory layers which decides the post-translational proteome of the parasites in particular disease conditions. Studies like these will help to decipher the post-translational environments of malaria parasites *in vivo* and elucidate their inherent proteome plasticity, thus allowing the conceptualization of novel strategies for interventions.

## BACKGROUND

The WHO World Malaria Report states that the global disease burden from malaria, caused by the protozoan parasites of the *Plasmodium* species, was 263 million cases in 2023 [1]. Deaths from malaria remain at an unacceptably high-rate post-pandemic [1]. Clinical manifestations of malaria include fever, chills, headache, myalgias, and fatigue. However, if not treated promptly, it may also lead to a severe disease state manifesting as jaundice, seizures, renal failure, acute respiratory distress syndrome, and coma. *Plasmodium falciparum* and *Plasmodium vivax* are primarily responsible for most of the disease burden. While the rapid progression of case morbidity in *P. falciparum* malaria had already been well-documented, an increasing number of reports of severe clinical complications caused by *P. vivax* infections have also been reported in the past two decades [2–7]. Recently, it has been reported that *Anopheles stephensi*, a mosquito vector capable of transmitting both *P. falciparum* and *P. vivax* has been detected in parts of the WHO African region and the incidence of *P.vivax* in these regions has increased from 0.5% to 1.1% in the last 5 years [1]. The recent reports of *P. vivax* resurgence in the USA [8], which had been certified malaria-free since the 1970s, remain a cause for concern.

In the human host, the parasite undergoes schizogony by invading RBCs involving a myriad of gene expression to help the parasite survive in the host cell [9]. Several studies have elucidated multiple mechanisms of post-transcriptional gene regulation in *Plasmodium* [9–13]. In addition to mRNA abundance and differential gene expression, epitranscriptomic studies in recent years have highlighted certain parasite-specific post-transcriptional modifications in *P. falciparum* as an additional layer of gene regulation [13,14,15]. A recent study highlighted that m5C modifications, particularly mediated by the NSUN2 methyltransferase, are vital for sexual differentiation and gametocyte production in *P. falciparum* and *P. yoelii* [16]. A knockout study of DNA/RNA methyltransferase, Pf-DNMT2 by Hammam et al., 2021 has revealed a substantial impact on parasite metabolism and stress response [17]. Another study has characterized the PfYTH2 reader protein in regulating the translational potential of m6A-modified mRNAs in *P. falciparum* [18]. However, as pointed out in the recent review by Rafael Serrano-Duran et. al.,2022 a major limitation of all the current epitranscriptomic studies in malaria is the lack of comprehensive simultaneous mapping of both m5C and m6A RNA modifications, particularly from ex-vivo studies involving patient-derived parasites [19]. The most commonly used platform i.e. LC-MS is conventionally used for detection of RNA modification, as employed in most of the previous studies [20, 21], however, in case of mixed human and parasite material as obtained from clinical cases it would be difficult to distinguish between human and parasite methylation signatures, in case of clinical samples, while, an appreciably greater amount of starting material would be required for antibody based assays which is rarely obtained in the field isolates. Long-read sequencing technology, such as from Oxford Nanopore Technologies can minimize these issues, enabling complete assembly of the parasite transcriptomes and cataloging of splicing events [21–22]. The current study attempts to address some of these concerns by elaborating on the RNA-specific methylation signatures along with their splicing patterns in malaria parasites by analyzing mono-infections of *P. vivax* and *P. falciparum,* as verified by the Polymerase chain reaction (PCR) assays. Both the clinical isolates show the same disease manifestation i.e. hepatic dysfunction (severe malaria).

The involvement of long non-coding RNAs and Natural Antisense Transcripts (NATs) has been demonstrated as crucial in mediating gene expression, thus influencing parasite development within human hosts [10,11,12,13,23]. Our previous work has profiled the abundance and differential expression of NATs in both *P. falciparum* and *P. vivax* patient isolates [10,11]. It may be hypothesized that methylation of the NATs can influence the stability and translational potential of the corresponding sense RNAs. Taking this into consideration, the current work also characterizes methylation signatures in NATs from the two species of the parasites under study; thus, adding a further complexity to possible post-transcriptional regulations. This is an aspect which has not been addressed earlier in parasite disease biology.

Alternative splicing (AS) is another mechanism that shapes the post-transcriptional landscape in eukaryotes, including *Plasmodium* spp., by generating transcriptome diversity and influencing protein structure and function [24]. It allows a single gene to produce multiple mRNA isoforms through selective exon recombination, governed by cis- and trans-acting factors [25]. AS is also likely to influence protein abundance through regulation of nonsense-mediated decay, and it is possible that some transcript isoforms that do not code for proteins have regulatory roles that are still unknown [26]. AS frequency varies across Apicomplexa, from 4.5% in *P. falciparum* to over 20% in *Toxoplasma gondii* [27, 28]. While its full functionality remains debated, studies suggest that AS plays a key role in transcriptional regulation, protein abundance, and parasite survival, making it a potential target for intervention strategies [29,30,31,32]. Although the short-read sequencing-based approach gives high depth to detect alternative splicing, it limits the identification of concurrent alternative splicing events, depicted by some apicomplexan gene splicing events that occur simultaneously on transcript isoforms [31]. The difficulty of identifying rare splicing events is further demonstrated by the absence of commonly used cutoffs or criteria for categorizing alternative splicing from RNA-seq [32]. However, long-read sequencing can minimize these issues, by transcriptome reconstruction, identifying novel isoforms and complex splicing events [27,32].

The genome of *P. falciparum* exhibits a high AT-content, of approximately 80%, one of the highest AT-rich genomes observed in any eukaryotic organism; significantly lower than *P. vivax*, which has a more balanced GC content of around 50% [33,34]. The genomic structure of *P. falciparum* is influenced by a strong mutational bias favoring transitions from guanine-cytosine (GC) to (AT) pairs. This results in a higher frequency of specific types of mutations, particularly G:C to A:T transitions, which can lead to rapid genetic changes that enhance the parasite’s adaptability to host immune responses and antimalarial drugs [35]. The AT-rich regions are often associated with genes involved in immune evasion, such as those coding for variant surface antigens. The concentration of these genes in sub-telomeric regions allows for rapid antigenic variation, which is crucial for the parasite’s survival in the human host [36]. In contrast, *P. vivax* has evolved a more stable genomic composition with a relatively higher GC content, contributing to genetic diversity. This suggests that it may experience different mutational pressures and evolutionary pathways compared to *P. falciparum*. The increase in GC content in *P. vivax* over time may be linked to mechanisms such as heteroduplex repair during meiotic recombination, which are less pronounced in *P. falciparum* due to its more clonal population structure [37]. It has already been suggested in literature, that the AT-GC contents of the respective parasites may give rise to different ways of influencing genome diversity. An added layer of methylation may provide a different perspective to understanding functional genome diversity.

This study elaborates on the epitranscriptomic landscape of *P. falciparum* and *P. vivax*, focusing on two RNA modifications - m5C and m6A, and their possible influence on post-transcriptional gene regulation. Using long-read sequencing, the current study provides single-base resolution of RNA methylation signatures in sense and Natural Antisense Transcripts (NATs), highlighting alternative splicing events. By addressing gaps in the comprehensive mapping of RNA modifications, particularly from patient-derived parasites showing hepatic dysfunction severe complication (PFC: *P. falciparum* complicated and PVC: *P. vivax* complicated), the work offers insights into parasite-specific regulatory mechanisms under the influence of various host factors. This preliminary data has the potential of indicating informed strategies for future therapeutic interventions.

## RESULTS

### Read statistics

Total RNA from two enriched, PCR-validated severe malaria samples was extracted as detailed in methods (Table S1-one each for *P. vivax* and *P. falciparum*). After quality and integrity check, the RNA was used for library preparation for Nanopore sequencing on MinION flowcell. In the case of *P. vivax*, we obtained 672k reads, while for *P. falciparum*, we obtained 207k reads. A dot plot of the read length vs. average read quality in Fig S1 depicts the mean read quality of both the runs ∼7. Only the base called reads above Q>7 (after basecalling by ONT basecaller Guppy v3.3.0) [38] were taken for further analysis. The longest mapped read length for PFC was ∼5 kb, while that for PVC was ∼3.5 kb. This shows the utility of long-read sequencing technology, performed for the first time in malaria patient-derived RNA samples. A detailed summary of the sequencing statistics is presented in Table S2.

### Transcriptome reconstruction and AS events

We further used the long read sequencing data to reconstruct the transcriptome, enabling the identification of alternative splicing events and novel genes using FLAIR and SQANTI3 [39, 40]. In total, 560,535 processed reads were analyzed, yielding 3,130 transcripts in the PVC dataset, derived from 1,582 annotated and 897 novel genes. These transcripts were further classified using SQANTI3 based on their structural similarities and differences compared to annotated transcripts. Among the identified transcripts, 1269 were full-splice matches (FSM) perfectly matching known transcripts, 50 & 335 were novel in catalog (NIC; refer to transcripts that contain new combinations of already annotated splice junctions or novel splice junctions formed from already annotated donors and acceptors) and incomplete-splice matches (ISM; refer to transcripts that matching part of a known transcript), respectively. Besides this, we identified a proportion of isoforms as antisense (NATs) (n=215), genic (n=362), intergenic (n=897), and fusion (n=2; transcriptional read-through between two or more adjacent genes) (Fig1).

**Fig 1.**
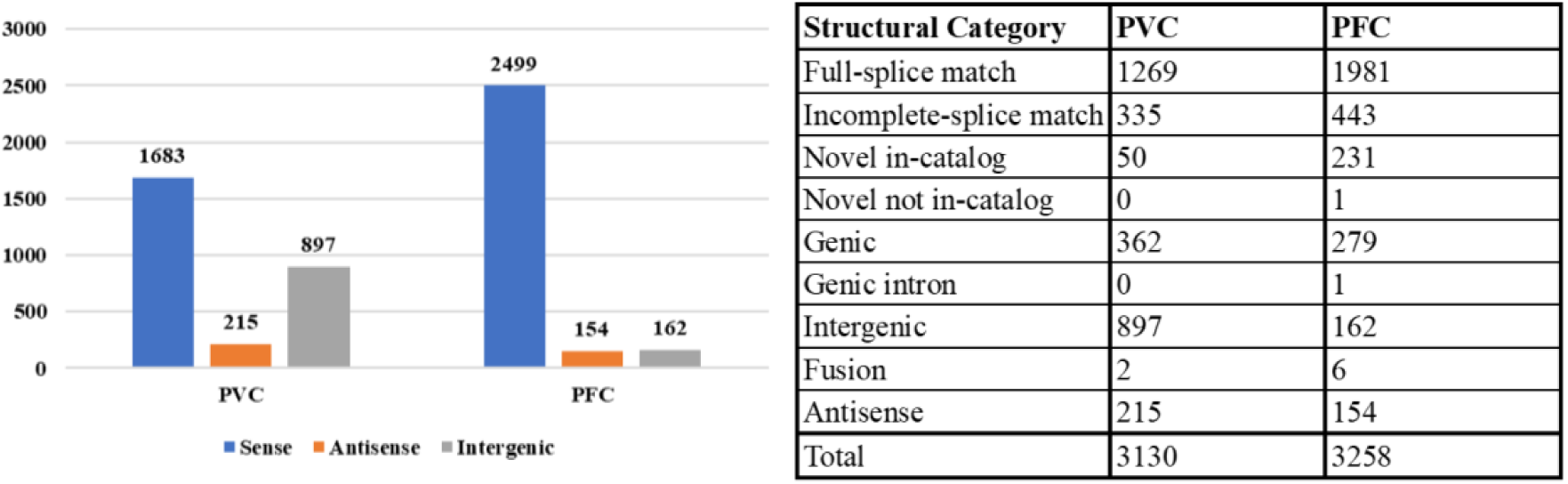
Number of isoforms expressed in PVC and PFC dataset segregated across the three (sense, antisense and intergenic) categories along with summary of transcript isoforms in PVC & PFC dataset based on structural categories of SQANTI3.

A total of 169,464 processed reads were analysed for the PFC dataset, leading to the reconstruction of 3258 transcript isoforms. SQANTI analysis provided a detailed classification of these isoforms spanning structural categories, with the majority being full-splice matches (n=1981). Additionally, a range of isoform categories were identified, including incomplete splice matches (n=443), genic (n=279), intergenic (n=162), antisense (NATs) (n=154), and novel in-catalog (NIC, n=231) (Fig1). Only one novel not-in-catalog(NNC) (New combinations of already annotated splice junctions or novel splice junctions formed with at least one novel splice junction) isoform (corresponding to gene PF3D7_0210100, 60S ribosomal protein L37ae, putative) was detected. Previously, this gene has been reported with 5’ and 3’ alternative splicing events [41]. We have detected this gene in the PFC dataset with “at least one novel splice-site” where the transcript length has reduced to approximately 50%. Six individual intra-chromosomal fusion isoforms and 1 genic intron isoform were observed, indicating the occurrence of a typical splicing events. The identified fusion transcripts exhibited either multi-exon splicing (n=3) or intron retention (n=3). The presence of multi-exon fusion transcripts suggests possible exon shuffling or novel gene fusions, while intron retention events may contribute to regulatory alterations, warranting further functional characterization. The most frequent alternative splicing event in both the datasets, identified was alternative 3’ end 5’ end splice sites (n=386 for PFC and n=336 for PVC).

Gene level annotation, after gffcompare [42], reports 834 expressed isoforms from PFC that are associated with novel transcripts and in case of PVC datasets, 1356 novel transcripts were reported, out of which 414 transcripts are from known genes.(Table-1) The detailed classification of all isoforms is provided in Supplementary Data 2 and 3 for PFC and PVC respectively.

**Table 1:**
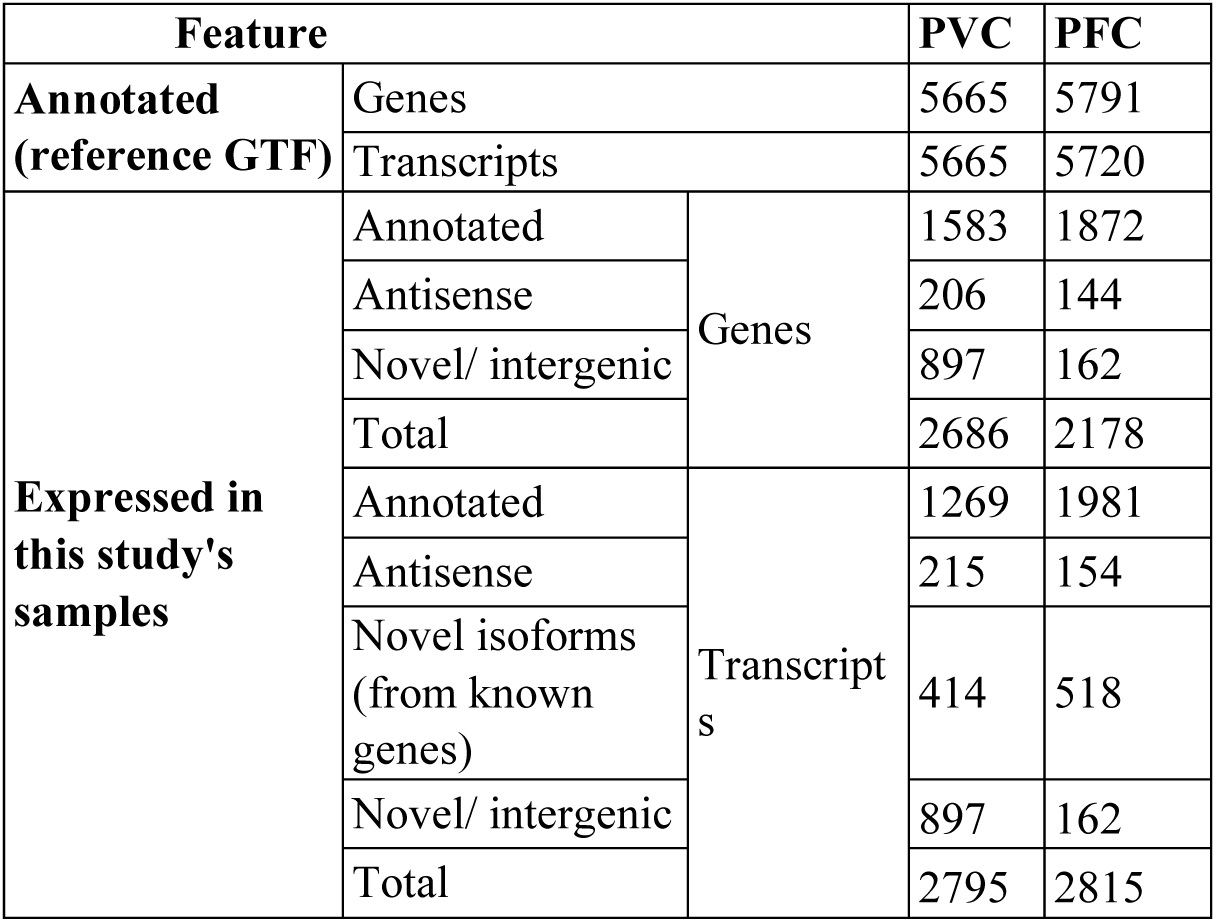
Summary of isoform count from FLAIR & SQANTI3 in PVC & PFC dataset.

| Feature |  |  | PVC | PFC |
| --- | --- | --- | --- | --- |
| Annotated<br>(reference GTF) | Genes |  | 5665 | 5791 |
|  | Transcripts |  | 5665 | 5720 |
| Expressed in<br>this study's<br>samples | Annotated | Genes | 1583 | 1872 |
|  | Antisense |  | 206 | 144 |
|  | Novel/ intergenic |  | 897 | 162 |
|  | Total |  | 2686 | 2178 |
|  | Annotated | Transcript<br>s | 1269 | 1981 |
|  | Antisense |  | 215 | 154 |
|  | Novel isoforms<br>(from known<br>genes) |  | 414 | 518 |
|  | Novel/ intergenic |  | 897 | 162 |
|  | Total |  | 2795 | 2815 |

### Distribution of methylation signatures across the genome

Methylation calling for reads reveals modifications in both sense, antisense and intergenic transcripts. Our data suggests RNA modifications of both m5C and m6A in all RNA types - mRNA, rRNA, ncRNA, snoRNA, and pseudogenic transcripts. All the m6A-modified transcripts detected by m6anet were covered in the modification calls predicted by CHEUI. Here, we report both the tools’ combined and concordant modification calls for m6A methylation. From the PFC sequencing run, a total of 824 m5C-modified and 836 m6A-modified unique isoforms were identified. Among these, 588 and 785 transcripts exhibited m5C and m6A modifications respectively with high confidence scores (≥0.9) (Fig 2e). While the co-occurrence of m6A and m5C modifications was found on 582 transcripts in the total dataset, no common methylated isoforms were observed between sense and antisense transcript pairs in PFC (Fig2d-f). Of the 588 m5C-methylated transcripts, 578 originated from sense transcripts, 3 were antisense (NATs), and 7 were intergenic (Fig 2d). Similarly, among the 785 m6A-modified transcripts, 767 were sense, 5 were antisense, and 13 were intergenic in origin (Fig 2f). We identified only one apicoplast encoded gene, PF3D7_API05900 (small subunit ribosomal RNA), as methylated in the PFC dataset. This gene, which encodes a non-coding RNA, harbored both m5C and m6A modifications at multiple locations. The presence of RNA modifications in both non-coding genes and NATs reinforces the idea that methylation is not exclusive to coding RNAs.

**Fig 2.**
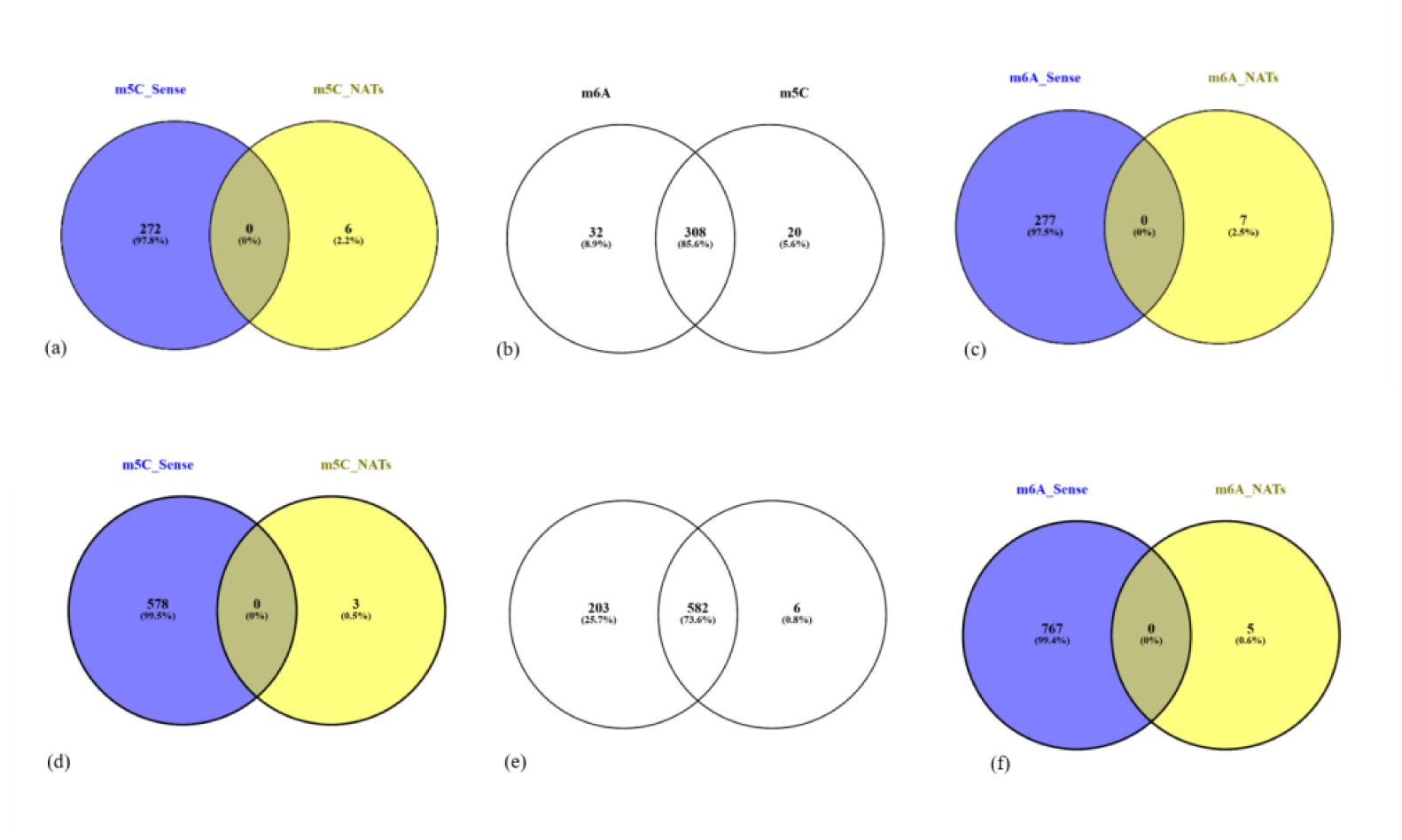
(a-c) Representation of genes in PVC data (from left to right) – m5C modification in sense & NATs, common transcripts between m6A & m5C modifications, m6A modification in sense & NATs, (d-f) Representation of genes in PFC data (from left to right) – m5C modification in sense, common transcripts between m6A & m5C modifications, m6A modification in sense.

In the PVC sequencing run, a total of 382 m5C-modified and 385 m6A-modified unique isoforms were identified. Among these, 328 and 340 transcripts exhibited m5C and m6A modifications respectively with high confidence scores (≥0.9) (Fig 2b). Of the 328 m5C-methylated transcripts, 272 originated from sense transcripts, 6 were antisense (Fig2a), and 50 were intergenic. Similarly, among the 340 m6A-modified transcripts, 277 were sense, 7 were antisense (Fig2c) and 56 were intergenic in origin.

In our analysis, 308 transcripts exhibited co-occurrence of both m6A and m5C modifications (Fig2b), out of which, several key genes and their functions, including mitochondrial transcripts, specifically targeting cytochrome c oxidase subunit I (PVAD80_MIT0002.1) and cytochrome c oxidase subunit III (PVAD80_MIT0001.1), which are critical components of the electron transport chain (ETC), which powers ATP production. Disruptions in their function can impair energy metabolism, forcing the parasite to rely on less efficient anaerobic pathways [43, 44]. Modifications in these genes could also increase resistance to antimalarial drugs like atovaquone by altering drug-binding sites or the overall functionality of the proteins [44,45]. Additionally, fusion genes (PVX_116525_PVX_116530) hypothetical proteins, conserved, (PVX_099320 & PVX_086090) glideosome associated protein & (PVX_090230) early transcribed membrane protein (ETRAMP), (PVX_111355) merozoite capping protein 1, putative, (PVX_096975)VIR protein,(PVX_111175) ookinete surface protein Pvs25, may enable the parasite to play multiple roles in regulating gene expression, protein abundance, and various cellular processes that are crucial for *Plasmodium* parasite survival, propagation and development throughout its complex life cycle [34, 46, 47, 48,49, 50].

### Mapping of methylation events to genomic features

Mapping of the transcripts showing RNA modifications to individual chromosomes for both PFC and PVC data was analyzed with respect to the event position and the GeneID. From this data, the overall distribution of these events across all the chromosomes appears to be even. However, we have not performed any enrichment analyses for the same. The liftover coordinates of methylation events in all isoforms are supplied in a bed file format in Supplementary Data 4.

The distribution of m5C and m6A modifications across sense, antisense, and intergenic transcripts in the PFC & PVC dataset, normalized by transcript length, show a clear increasing trend along the transcript length in the sense isoforms, with the highest frequency near the 3’ end (normalized length close to 1.0) (Fig 3). Further mapping of methylated sense isoforms with full-splice matches in the PFC dataset to the genomic features revealed the positions of 444 methylated transcripts mapped to 3’UTR, which is the highest in comparison to 367 and 67 modified bases mapped to CDS and 5’UTR regions of the mRNA transcripts respectively. For m5C modification detected for a total of 376 mRNA transcripts, 277 methylation events lay in the coding segment, while 35 and 255 m5C-methylation base fell in the 5’UTR and 3’UTR region, respectively. The Venn diagram shows the clustering of RNA modifications in more than one region of the transcript, where 18 transcripts in the PFC dataset have m5C methylated site lying in all the three - 5’UTR, CDS, and 3’UTR regions of the mRNA, whereas 58 transcripts show m6A methylated bases spread across all three annotated regions of the mRNA (Fig 4).

**Fig 3.**
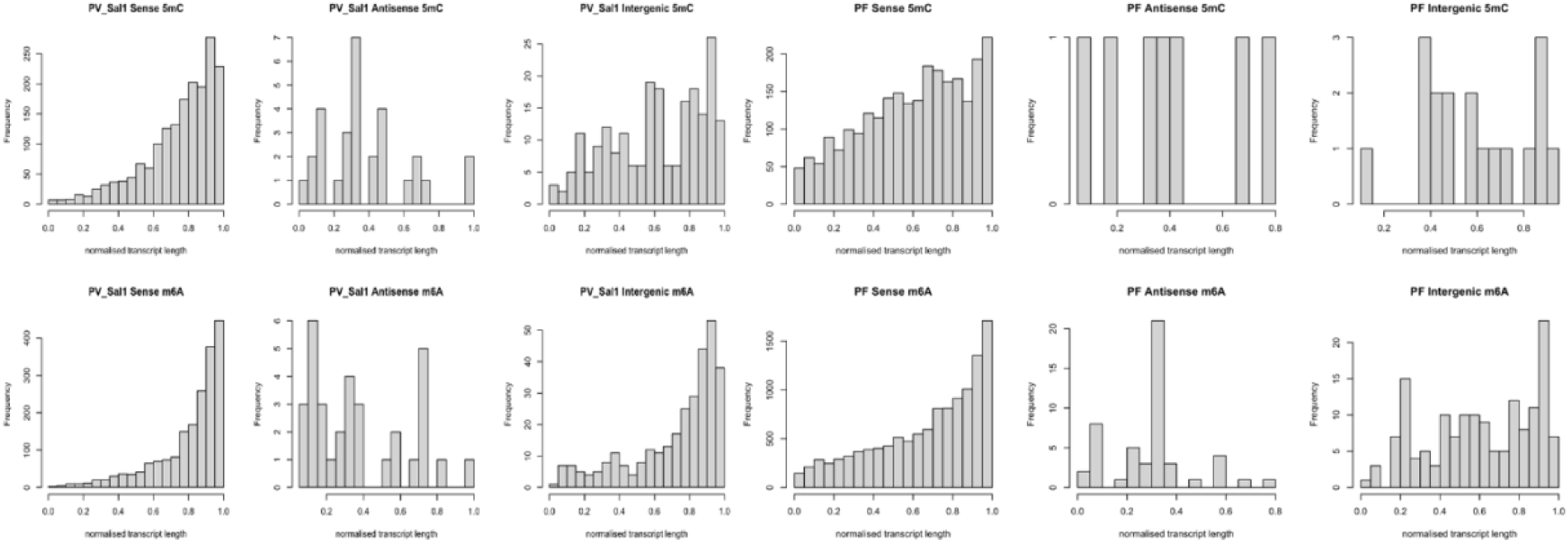
Histogram representation of modified base distribution across normalized transcript length.

**Fig 4.**
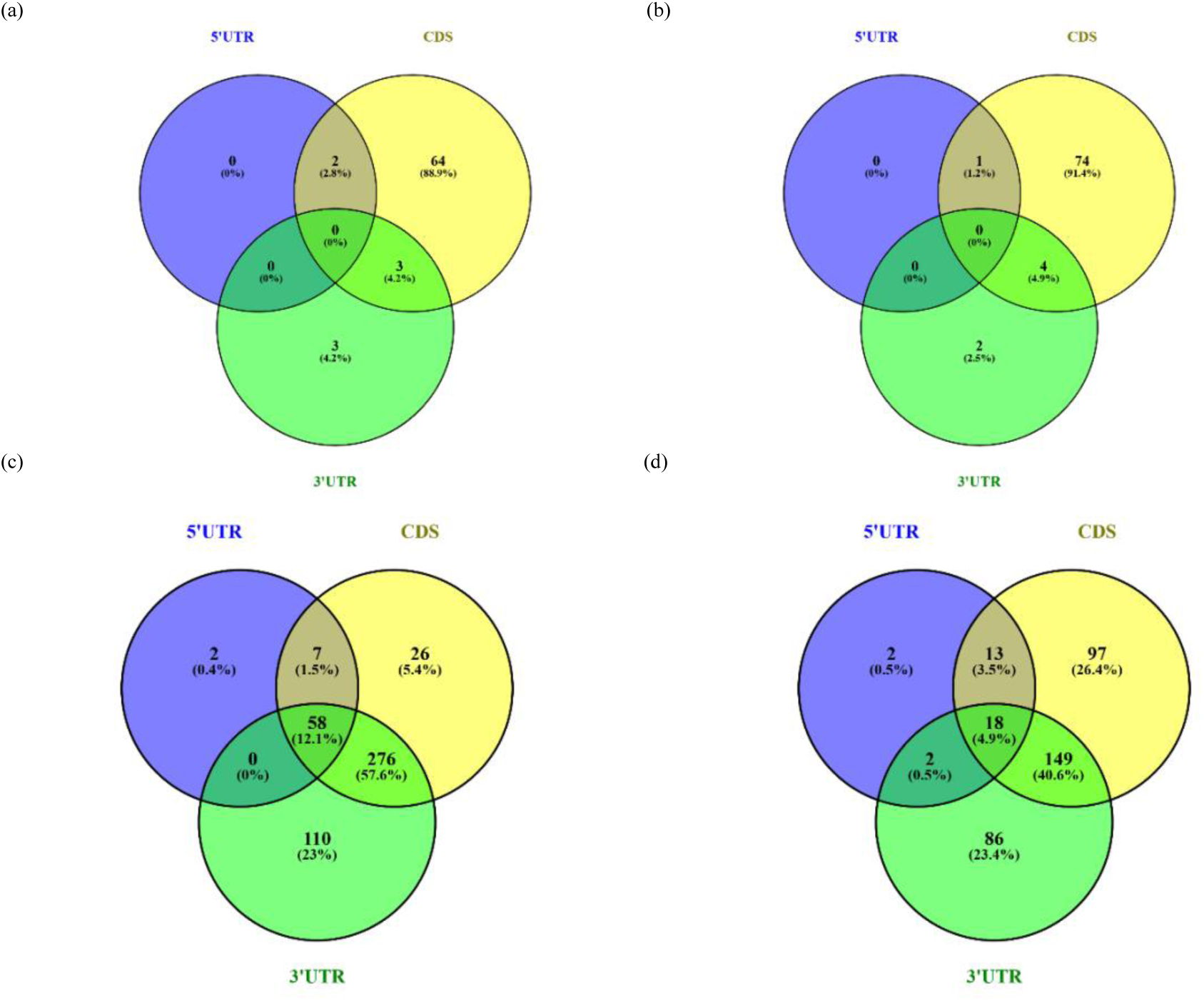
**(a)**-Number of FSM mRNA transcripts with methylation events (m6A), which mapped to different annotated mRNA features in PVC. **(b)** Number of FSM mRNA transcripts with methylation events (m5C), which mapped to different annotated mRNA features in PVC. **(c)** Number of FSM mRNA transcripts with methylation events (m6A), which mapped to different annotated mRNA features in PFC. **(d)** Number of FSM mRNA transcripts with methylation events (m5C), which mapped to different annotated mRNA features in PFC. *Detailed transcript IDs are provided in Supplementary Sheet 3.

In case of PVC datasets, a total of 199 mRNA transcripts with one or more m6A modification sites are detected, methylation events of 69 mRNA transcripts in the PVC dataset are mapped to CDS, which is the highest in comparison to 6 and 2 transcripts for 3’UTR and 5’UTR regions, respectively(Fig 4).For m5C modification detected for a total of 194 mRNA transcripts, methylation events of 79 transcripts lay in the CDS, while 1 and 6 m5C-methylated transcripts fell in the 5’UTR and 3’UTR region, respectively (Fig 4).The Venn diagram shows the clustering of RNA modifications in more than one region of the transcript, in m5C. One of these transcripts(PVX_111175.1-ookinete surface protein Pvs25) has m5C in the 5’UTR & CDS. The other four transcripts (PVX_114910.1-40S ribosomal protein S15, putative, PVX_098655.1-hypothetical protein, conserved, PVX_087960.1-inner membrane complex protein 1d, putative, PVX_080520.1-hypothetical protein, conserved) have m5C modifications in the 3’UTR & CDS regions of the mRNA. In m6A, two transcripts (PVX_111175.1-ookinete surface protein Pvs25, PVX_118115.1-hypothetical protein) have modifications in the 5’UTR & CDS and three transcripts (PVX_087960.1-inner membrane complex protein 1d, putative, PVX_114910.1-40S ribosomal protein S15, putative, PVX_098655.1-hypothetical protein, conserved) have modifications in the 3’UTR & CDS regions of the mRNA.

### Gene Ontology Enrichment Analysis

Functional enrichment analyses were performed individually for the m5C and m6A methylated transcripts (detected with a high confidence modification probability score of >=0.9) using Gene Ontology (GO), and KEGG databases to interpret the biological relevance of the modification events. Across the two datasets, genes for both m5C and m6A modifications were reported to be enriched for biological processes corresponding to translation (GO:0006412), gene expression (GO:0010467), cytosolic large (GO:0022625) and small (GO:0022627) ribosomal subunit. Apart from this, individual GO terms are identified separately in PFC and PVC datasets (Supplementary Data 5).

In both the datasets, the enrichment of genes in the KEGG pathway revealed – ‘Ribosome (pfa03010/pvx03010),’ along with the genes associated with the biological process of translation. Among the significant KEGG pathways identified in the PFC methylation dataset are - pfa00010:Glycolysis/Gluconeogenesis and pfa01230:Biosynthesis of amino acids, associated with biological processes like – carbohydrate catabolic process (GO:0016052), ribonucleotide catabolic process (GO:0009261), and glycolytic process (GO:0006096), suggesting activation of pathways involved in cellular energy production and nucleotide metabolism. This suggests that RNA modifications in *P. falciparum* might alter the metabolic adaptations in the parasite in response to its host cell environment, specially in the case of severe disease.

Apart from these house-keeping metabolic and biogenesis processes, the gene ontology enrichment analysis of methylated m6A and m5C sense transcripts in the PFC dataset has also identified highly specific biological processes associated with translation, ribosome biogenesis, and RNA metabolism (GO:0006412, GO:0006414, GO:0001732, GO:0022618 and GO:0019843). These findings indicate the potential involvement of methylated transcripts in fine-tuning translational efficiency in the parasite. Significant enrichment was observed in vesicle-mediated transport pathways, including extracellular (GO:1903561) and intracellular vesicle (GO:0097708), phagocytic vesicle (GO:0045335), lysosome (GO:0005764) and translocation of proteins into host (GO:0042000), suggesting that m6A modifications may also influence intracellular trafficking, possibly aiding in nutrient acquisition. The enrichment of GO:0036396 - RNA N6-methyladenosine methyltransferase complex exclusively in the m6A methylated PFC dataset identifies transcripts directly associated with the m6A modification machinery and leads to interesting possibility of whether m6A modifications help regulate their own deposition in a feedback mechanism.This could be a lead to further investigations.

The gene ontology (GO) for *P. vivax*’s unique sense transcripts exhibiting m5C and m6A modifications (as detailed in Supplementary Data 5) highlights the involvement of genes in two significant biological processes: cytoskeleton organization (GO:0007010) and protein-DNA complex organization (GO:0071824). These processes are crucial in governing the parasite’s shape, invasion strategies, and gene expression patterns. Additionally, the presence of unique antisense transcripts associated with RNA methyltransferase (GO:0008173) and S-adenosylmethionine-dependent methyltransferase (GO:0008757) functions suggests the existence of an intricate regulatory system. This complex network is likely responsible for precisely modulating gene expression throughout the various stages of the parasite’s life cycle, potentially enabling it to circumvent host immune defenses and adjust to diverse cellular environments.

### Orthologs in the two datasets

We further identified orthologs of methylated sense transcripts (43 for m6A and 37 for m5C modification) across *P. falciparum* and *P. vivax* data, revealing conserved epitranscriptome patterns (Supplementary Data 6). This cross-species analysis hints at the potential evolutionary importance of methylation in *Plasmodium,* which may impact translational repression or activation in severe malaria [51]. One class of molecules exhibiting m6A methylation are some members of the protease family (PF3D7_1115300, cysteine proteinase falcipain 2b and PVX_091405, vivapain-2). These molecules have been well characterized as a potential drug target against malaria [52]. It is important to consider the modification of these transcripts, which may influence the translation potential of these key molecules. Transcripts from another pair of orthologous genes PVX_090230 and PF3D7_0202500 encodes for early transcribed membrane protein 2 (ETRAMP2) and displays m6A modification on sense transcripts of both the datasets with a high confidence of 0.999. Showing differential expression in severe cases of malaria with a high frequency of m6A modification, ETRAMP2 could be an important antimalarial target requiring further investigation. In the case of m5C, most modifications in the orthologs of the two datasets are seen in transcripts for ribosomal proteins suggesting that these may be stabilizing the translational machinery.

### Comparison with existing datasets of RNA modification

A comparison between the RNA modification datasets of *P. falciparum* cultured strains and the clinical isolates sequenced in the current study highlights intriguing overlaps despite substantial differences in experimental conditions and biological sources. Analysis of m5C-modified transcripts in *P. falciparum* 3D7 across different developmental stages analysed by Liu et al., 2022 [16] using Bisulfite sequencing revealed overlap of only 3% of methylated genes in the schizont stage, while 5.7%, 3%, and 4.4% were identified in gametocyte stages III, IV, and V, respectively. When pooling transcripts (with at least one m5C site) across all stages and comparing them with the PFC dataset, only five transcripts were consistently methylated. These five common transcripts included three ribosomal RNAs (PF3D7_0532000, PF3D7_0725600 and PF3D7_0726000), a conserved *Plasmodium* protein of unknown function (PF3D7_1135700), and a putative elongation factor 1-gamma (PF3D7_1338300). Their presence across multiple stages of culture and blood-stages from clinical setting, suggests a potential role for m5C modifications in translational regulation and highlights the importance of ribosomal RNA methylation in parasite gene expression. Similar investigation of m6A modification dataset of PFC with m6A-enriched peaks reported by Baumgarten et al., 2019 [20], revealed stage-specific overlaps, with 9.1%, 8.9%, and 7.9% of methylated genes detected at 12-, 24-, and 36-hours post-infection (h.p.i.), respectively. When pooling all time points, 147 genes (10%) were found to be commonly methylated between the two m6A modification datasets (Supplementary data 7).

Comparison of the PVC dataset with the published datasets showed orthologs identified for *P. vivax* Sal 1, with variable levels of overlap across the m6A and m5C datasets (Supplementary data 7).

### Differential methylation of isoforms

As long reads ideally capture the entire transcript, reads with RNA modification must correspond to transcript molecules with that modification. Therefore, we tested the isoforms detected for specific methylation, i.e., some isoforms of a gene preferentially undergoing methylation (m6A and/or 5mC) versus others. Applying Fishers’ test to the genes with m6A and/or 5mC RNA modifications detected in either species with at least two isoforms were tested for differential RNA methylation of isoforms (Fig S2). This yielded only one gene PF3D7_0913300(conserved protein, unknown function) in the PFC dataset with its major isoform preferentially methylated for both m5C and m6A modifications. Since this isoform is associated with a novel transcript of the gene, the mapping of methylation coordinates for its transcript could not be attempted. However, in the case of PVC, there were no significant differentially methylated transcripts (Supplementary Data 8).

## DISCUSSION

More than 130 chemical modifications of RNA are known to date [51], but out of these m6A & m5C are the most prevalent modifications studied. m6A is one of the most abundant internal modifications on mRNA, affecting RNAs’ splicing, translation, stability, and nuclear export. *P. falciparum* exhibits relatively high m6A deposition levels, which surpass those observed in human mRNA. This is attributed to the parasite’s adenosine-rich protein-coding transcriptome, suggesting a key role for m6A modification in regulating its asexual replication cycle [20]. They are strictly regulated by methyltransferase (MTC) writer complex (PfMT-A70) identified by the YTH reader homolog [18]. Building on this foundational understanding, the current study expands the scope by investigating RNA modifications in clinical isolates across two *Plasmodium* species. While m6A modifications show consistent patterns of deposition with that of m5C in *P. falciparum*, the relatively low overlap of this RNA modification with the m6A dataset published by Baumgarten et. al. 2019 [20] may be attributed to the ex-vivo sequencing of patient isolates in our study as opposed to the highly synchronous *P. falciparum* 3D7 culture used by Baumgarten et. al [20]. As for comparison with the dataset of mRNA-BisSeq by Liu et al., 2022 [16], only five sense transcripts modified for m5C are common between PFC dataset and all parasite stages studied by the authors. Several confounding factors like the stage-specific regulation of RNA modifications, parasite’s diverse genetic backgrounds and differences in experimental approach may limit the concordance of RNA modification sites reported by earlier studies. However, the majority of m6A sequencing reads in the cited study match to regions that code for proteins [14, 20, 53]. Our study also supports the previous findings for m6A as well as m5C since most modifications are present in the coding segments of the mRNA. Similarly, in *P. vivax* isolates, comparable patterns were observed, with a low overlap in RNA modification sites in *P. vivax* Sal-1 which were not surprising, given the clinical nature of the isolates and the inherent differences between the species.

None of the studies cited have investigated the occurrence of both m5C and m6A modifications in the same transcripts. The presence of both of these modifications on 73% of transcripts (Fig 2e) in PFC and ∼85% of transcripts in PVC dataset (Fig. 2b) highlights a possible intricate mechanism by which *Plasmodium* fine-tunes its transcriptome in response to environmental changes, optimizing survival and pathogenicity within the host and evading immune responses [14,16,19,53,54]. The distribution of both types of modifications in both the datasets is predominately much lower in the 5’UTR segment of the transcripts. This strong preference of methylation in the coding segment of the gene may suggest CDS to be the primary target for gene regulation mediated by RNA modifications rather than the UTRs. These dual modifications could provide a sophisticated regulatory framework that may govern transcript stability, localization, and translation, ultimately shaping cellular responses and functions. The tandem analysis of methylation patterns from transcripts originating from orthologous genes is important to understand the potential implications of these gene products during the process of disease severity. Transcripts of the ETRAMP family are orthologous in both datasets (Supplementary sheet 4) with very stringent methylation probabilities dispersed at distinct annotated regions of mRNA. Proteins of the ETRAMP family are known to be widely expressed in the early asexual stages of all *Plasmodium* species at the host-cell interface [48,49]. Upon early intra-erythrocytic development, the parasites need to export much of their proteins into the host cell to sustain their survival within the host. This molecule has an important role in transporting proteins outward from the parasite, and this role could be enhanced in cases of severe disease [48].

Hemoglobin hydrolysis, invasion, and egress from host erythrocytes are all impacted by cysteine proteases of the papain-family [55, 56]. These molecules have been characterized as chemotherapeutic targets [56]. m6A methylation detected in orthologous transcripts encoding for members of this family may result in altered protein expression profiles, which could challenge their role as potential chemo-therapeutic targets [56]. This knowledge is invaluable for identifying new targets for therapeutics, particularly those that are indispensable for the parasite’s survival, thereby supporting the development of more effective interventions against malaria.

In addition to base modifications, alternative splicing provides an evolutionary mechanism to generate a complex proteome from a relatively limited number of genes. In *P. falciparum*, with a relatively dense genome, even modest alternative splicing events can critically modulate protein expression and influence host–parasite interactions, potentially affecting parasite virulence and immune evasion [57]. They have reported high numbers of spliced isoforms in the members of the Ap2 transcription factor family. This illustrates the importance of alternative splicing in modulating Ap2-mediated gene regulation in *Plasmodium* [57]. Comparative studies in more complex eukaryotes further illustrate that precise splicing regulation is integral not only to neuronal plasticity and signal transduction [58] but also to broader evolutionary adaptations that enhance cellular functionality [59]. Splicing of various sense and antisense transcripts with m5C and m6A modification events has been seen in our datasets. Despite *P. falciparum* being a well-annotated genome, we report new transcripts and splicing events not present in the existing annotations of the PF3D7 genome(v64) from PlasmoDB [60]. Our splicing data in PFC reports 6 individual fusion events. All the six fusion events are intra-chromosomal (Supplementary Data 2) in comparison to the same transcripts reported by the study of Yang et. al.,2021 as inter-chromosomal, using PacBio long read sequencing on *P. falciparum* 3D7 [57]. Both these studies are an important first step in characterizing the clinical implications of fusion events in malaria and warrants further investigation with respect to occurance of chimeric proteins.

*P. vivax* exhibits alternative splicing similar to *P. falciparum* [41, 61]. We have found two intrachromosomal fusion events in PVC dataset (Supplementary Data 3) i.e. hypothetical proteins with unknown functions-PVX_002870; PVX_002867 & PVX_116525; PVX_116530, which may indicate a major role of splicing in influencing proteins with potentially altered activity. It is noteworthy that in our analysis, 50 isoforms, which were cataloged as Novel in-catalog, exhibited diverse splicing patterns when compared with previously published data by Bozdech et al., 2016 [61]. Specifically, several genes demonstrated alternative splicing events, including combinations of known splice sites, alternative 3’ and 5’ ends, 3’ prime fragments, and intron retention. For instance, PVX_118115, PVX_113790, and PVX_099015 were identified as having a combination of known splice sites, aligning with AS5 or AS3 categories in Bozdech’s data.,2016. PVX_084260 exhibited an alternative 3’ and 5’ end, corresponding to the AS5 classification. Additionally, PVX_091395 presented as a 3’ prime fragment, also categorized as AS5. Interestingly, PVX_097875 and PVX_079725 predominantly exhibited intron retention, a feature that was classified under AS3 in some instances, whereas in other cases, Bozdech et al.,2016 categorized them as exon skipping. This variation suggests potential differences in transcript annotation or regulatory mechanisms affecting these genes.

Apart from this, there were also other instances of intron retention (IR) (Supplementary Data-3), with some predicted to undergo non-mediated decay (NMD) while others do not. According to the literature, intron retention is frequently associated with the downregulation of gene expression via NMD (IR-NMD) [62] as retained intronic sequences that disrupt the main open reading frame (ORF) often introduce premature termination codons (PTCs). The fate of an mRNA containing one or more IR events is influenced by various factors, including the position of the retained intron within the transcript. Overall, IR can affect ribosomal association, either by acting as a precursor to NMD or by modulating translation initiation through upstream open reading frames (uORFs) or other regulatory sequences that either repress or enhance translation initiation [63] Based on our data, this could suggest that IR serves as an important component of gene expression regulation via post-transcriptional splicing of retained introns during severe malaria.

Overall, these observations highlight the complexity of alternative splicing phenomenon that expands the proteomic diversity by generating multiple mRNA isoforms from a single gene, thereby fine-tuning protein functions and supports its exploration as a potential therapeutic target in infectious diseases [57, 64]. This may imply malaria manifestation driven splicing events by the parasite and should be further investigated by increasing the sample pool. Integration of this isoform data with other proteomics/metabolomics data can contribute to a comprehensive understanding of molecular networks underlying malaria/severe or complicated malaria.

The methylation events (m6A & m5C) in PVC dataset are predominantly concentrated in the coding regions of sense transcripts, followed by the 3’ and 5’ UTRs. It has been earlier reported [18] that m6A mRNA modification in the CDS reduces the stability of the transcripts or takes it into a degradative pathway. Other studies have shown that modification at the 3’UTR regions enhance RNA degradation [65]. Our data shows the presence of significant amounts of m6A modifications predominantly in the CDS of the mRNA transcripts (Figure 4a). This could suggest enhanced mRNA degradation, potentially decreasing translational efficiency of some transcripts in case of severe disease. Conversely, m5C modifications are associated with translational fidelity and subcellular localization. Enriched in coding regions, particularly downstream of the translational start site, m5C modifications enhance translation initiation [66]. The interplay between m6A and m5C possibly fine-tunes the translational landscape, enabling the parasite to adapt to environmental cues and stress conditions [66,67]. Both modifications also regulate RNA stability and degradation. While m6A can promote rapid transcript decay by recruiting decay machinery, m5C may stabilize transcripts by preventing degradation [65, 66]. Apart from these, our data also highlights previously unannotated transcripts in *Plasmodium* emphasizing the importance of ex-vivo transcriptomic studies in malaria research.

It has been earlier reported [20] that m6A mRNA modification in the CDS reduces the stability of the transcripts or takes it into a degradative pathway. Other studies have shown that modification at the 3’UTR regions enhances RNA degradation [65]. Our data shows the presence of only m6A in 8.9% & 25.7% transcripts in PVC & PFC datasets respectively (Fig 2b,e). In these m6A modifications are seen predominantly in the CDS or 3’UTR of the mRNA transcripts (Figure 4a,c). This could suggest enhanced mRNA degradation, potentially decreasing translational efficiency of some specific transcripts in case of severe disease which could potentially affect parasite survivability under these conditions. Additionally, the GO analysis underscores the reliance of *P. vivax* on significant biological process for its survival and highlights potential targets for antimalarial intervention.

## CONCLUSION

This is the first report to comprehensively profile the m5C and m6A modification distribution in sense and NATs from the clinical transcriptome of *P. falciparum* and *P. vivax*. These modifications may potentially modify mRNA stability and translation. This study faces several challenges inherent to working with clinical *Plasmodium* isolates and nanopore sequencing. The absence of replicates further complicates differential analyses. Additionally, low sequencing coverage and human RNA contamination reduce data resolution. Despite these limitations, our study provides several facets of post-transcriptomic mechanisms operating in severe malaria patient-derived *Plasmodium* isolates of both *P. falciparum* and *P.vivax*. Further, we have identified methylated transcripts encoded from *Plasmodium*’s nuclear, mitochondrial and apicoplast genomes. A detailed analysis of the quantification of splicing events in methylated sense and NATs is presented. This study reports several conserved hypothetical genes and novel transcripts from the two *Plasmodium* species, opening future avenues for investigating and characterizing these new molecules. Our study stresses the need to sequence the parasites in their natural host environment and lays the foundation for future functional validation, emphasizing the need for both *in vitro* and *in vivo* studies to validate these post-transcriptomic findings and their relevance to parasite survival and pathogenesis.

## METHODS

### A. Sample Collection and processing

Venous blood samples (5ml) from patients infected with malaria were collected from S.P. Medical College, Bikaner, with the informed consent of the patients according to the guidelines of the relevant ethics committee. Malaria infection with *P. vivax* and *P. falciparum* was confirmed with RDTs (OptiMAL, Flow Inc., Portland, Oreg.) and microscopic investigation of peripheral blood films. The blood samples were separated on a Histopaque-gradient-based density separation (Histopaque 1077, Sigma Aldrich) to separate the erythrocytes from plasma and peripheral blood mononuclear cells (PBMCs). The erythrocytes were washed with sterile PBS, lysed with TRI Reagent (T9424, Sigma-Aldrich), and transported in a cold chain to BITS Pilani for further processing.

Total RNA was isolated from infected erythrocytes according to the manufacturer’s protocol (Tri-Reagent, Sigma Aldrich) and resuspended in nuclease-free water. Molecular confirmation of infection with P. vivax or P. falciparum was done using 18S rRNA gene-based multiplex PCR and 28S rRNA gene-based nested PCR [68, 69], and any possibility of mixed infection was negated. Viral or other causes of hepatic dysfunction were ruled out at the point of collection. This study used one sample each of *P. vivax* and *P. falciparum*. The patient’s diagnosis was confirmed as hepatic dysfunction as per WHO guidelines [70] and upon our team of clinicians’ consultation. The clinical characteristics of the sample are tabulated in Table S1.

### B. Library Preparation

The concentration and purity of the isolated total RNA were measured by the NanoDrop UV/Vis spectrophotometer (SimpliNano™ spectrophotometer, GEHealthcare) and Qubit 4 Fluorometer (Qubit RNA HS Assay kit (Q32855), Invitrogen). The quality of the isolated total RNA was analyzed by denaturing agarose gel. One µg of total RNA was used for library preparation as per the manufacturer’s protocol (SQK-RNA002, Oxford Nanopore Technologies), including the optional reverse transcription step recommended by ONT. ∼20 ng of reverse transcribed and adapter-ligated RNA was used for loading on Spot-ON Flow Cell (R9) and sequenced on MinION sequencer (FLO-MN106D, Oxford Nanopore Technologies). Each of the two samples was sequenced on individual flow cells with repeated loading of 20 ng of the final library until exhaustion of pores, which gave a run time >=24 hours.

### C. Transcriptome reconstruction

Reads generated at the end of the sequencing procedure were basecalled using the base caller integrated into the MinKNOW software (v23.07.15; Guppy basecaller v3.3.0) [38]. After base calling, the reads categorized in the “pass” folder were concatenated and used for downstream analysis. ONT sequencing data was checked for quality using FastQCv0.12.1. The reads were indexed with Nanopolishv0.14.0 (https://github.com/jts/nanopolish) [71] using preset commands and mapped to the respective genomes (*P. falciparum* 3D7 for PFC and *P. vivax* Sal1 for PVC dataset) using minimap2 [72] *-ax splice* parameter. Aligned reads were corrected at splice junctions, collapsed into unique isoforms, quantified by FLAIR (Full Length Alternative Isoform Analysis of RNA) [39] using default parameters and some custom commands; followed by comparison with reference annotation using gffcompare [42]. A read-to-transcript map was also generated at the collapse step to filter the methylated vs non-methylated reads identified downstream (see section D). This was followed by transcript classification into full-splice match (FSM), incomplete-splice match (ISM), antisense (AS), novel in-catalog (NIC), novel not-in-catalog (NNC), genic (G), and intergenic (IG) based on similarity to existing annotations, using SQANTI3 [41]. For all downstream analysis, ISMs were removed. We have sub-divided all expressed isoforms into three bins as per the SQANTI categorization, corresponding to – sense (Full splice match-FSM, genic, genic intron, fusion, novel-in-catalog-NIC, novel-not-in catalog-NNC), antisense and intergenic isoforms. These “antisense” transcripts, as described by the tool [41], represent the NATs, which we have earlier reported [10, 11]

### D. Methylation Calling and downstream bioinformatics analysis

Methylation calling in sense and antisense isoform bins (as described above) was done by m6anet (https://github.com/GoekeLab/m6anet) for m6A modification calls and CHEUI: Methylation (CH3) Estimation Using Ionic current (https://github.com/comprna/CHEUI) for both m5C and m6A modification calls [73,74]. The cumulative modification calls from the two tools were considered for further analysis. The analysis considered only the modified calls with a read_coverage of ≥20 and modified_probability >=0.9 to maintain data stringency. The genes that satisfied the methylation criteria were used for functional enrichment analysis by DAVID (https://david.ncifcrf.gov/) with a p-value cut-off (EASE score) <0.1 against a background list of all expressed genes in the individual sequencing data. Transcripts showing methylation with the above-mentioned confidence were mapped to their position on the individual chromosome using lift-over script presented in R2Dtool [75]. For all mRNA transcripts that were full-splice match to the reference (and corresponded to the annotated features), the modified base position was mapped to genomic features as per the annotation of the reference genome using a custom R script.

### Differential methylation of isoforms

A single list of methylated readIDs identified from CHEUI and/or m6anet for 5mC and m6A methylation marks were created per species. These were then used to filter the read-to-transcript map generated with FLAIR to obtain the set of methylated transcripts, in R. Genes with at least two isoforms and at least one methylated read (of either mark) was filtered and major isoform was determined based on counts. A 2×2 contingency table was made with (a) all methylated counts of the gene, (b) methylated counts of the major isoform, (c) total counts of the gene, and (d) total counts of the major isoform (Fig S2). Fisher’s test was performed, and log odds ratio (LOR) was calculated as 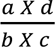 for each tested gene. A negative LOR indicates more methylated reads detected for the major isoform as compared to that for the gene and vice-versa. The p-values obtained from Fisher’s test were corrected for multiple testing using Benjamini-Hochberg (BH) procedure. All associated codes are uploaded to GitHub (https://github.com/pgupta3005/Plasmodium_LR) and intermediate files for all isoforms used in this analysis and their counts used as inputs in the contingency tables are given in Supplementary data 8.

## DECLARATIONS

### Ethics Approval

Sample collection was earlier approved by SP Medical College Hospital’s Ethics Committee (No.F. (Acad)SPMC/2003/2395). Permission to use these samples for further studies was given through IERB approval No. F29(Acad)SPMC/2020/3151 dated 05.09.2020. We thank all the patients for their voluntary consent and participation.

### Consent for publication

Not applicable

### Availability of data and materials

The direct RNA Sequencing raw data generated in this study have been deposited in the Indian Nucleotide Data Archive (Accession #INRP000148) and INSDC (Accession #PRJEB75779). The direct RNA-Seq data are available under controlled access to ensure strict confidentiality. Access can be obtained by submitting a request to our Data Access Committee. All the sequencing data used for this study will be available immediately after publication. All data generated are provided in the main text and supplementary data files. All patient meta-data included in this article are included in Table S1. Reprint requests should be addressed to AD. All Venn diagram illustrations have been made using Venny [76] while histogram have been made in R.

### Competing Interests

All authors declare no competing interests.

### Funding

This work is funded by the Indian Council of Medical Research (ICMR) under the Extramural Ad-hoc Scheme (PID: 2019-1121).

### Author Contributions

AD conceived the study, organized the material, was the principal investigator of a project from ICMR, India, and networked to facilitate the outcome. SG and PR were involved in performing the sequencing experimentation and subsequent analysis. AD, SG, and PR wrote the manuscript. SG and PR are equal contributors to this work. PG and IG performed transcriptome reconstruction and analyzed the differentially methylated transcripts. IG has been closely involved in discussions since the study’s inception. TM supervised the application of various in-silico tools and assisted in troubleshooting when required. DG was involved in consultations on data analysis and related discussions. DKK and SKK supervised sample collection and clinical characterization. All authors read and approved the manuscript.

## Acknowledgments

This work is funded by the Indian Council of Medical Research (ICMR) under the Extramural Ad-hoc Scheme (PID: 2019-1121) to AD, Department of Biotechnology, Government of India, through the Ramalingaswami Fellowship (ST/HRD/35/02/200) to IG and Ministry of Human Resource Development, Government of India sponsored Prime Minister’s Research Fellowship (PMRF) (IITD/Admission/PhD/PMRF/2020–21/380485) to P.G. We thank Birla Institute of Technology (BITS), Pilani (Pilani campus), International Centre for Genetic Engineering and Biotechnology (ICGEB), New Delhi, Indian Institute of Technology (IIT), New Delhi, India and SP Medical College, Bikaner, for providing the necessary facilities required for this work. SG and PR were supported by ICMR as project assistants to carry out this work and were subsequently supported by Institute fellowships for the PhD program applicable by BITS Pilani (Pilani campus).

## Notes

### Competing Interest Statement

The authors have declared no competing interest.

### Summary of Updates

We have revised the analysis according to the reviewer comments.

