## Supplementary_File for "A Tale of Two Parasites: A Glimpse into the RNA Methylome of Patient-derived *Plasmodium falciparum* and *Plasmodium vivax* isolates": SupplementaryData1.docx

**Supplementary Figures**

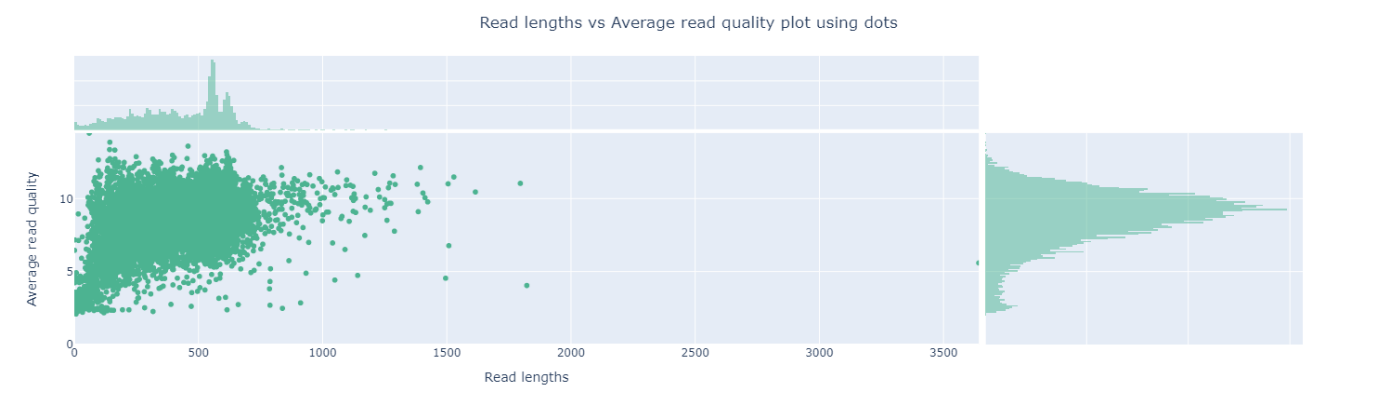

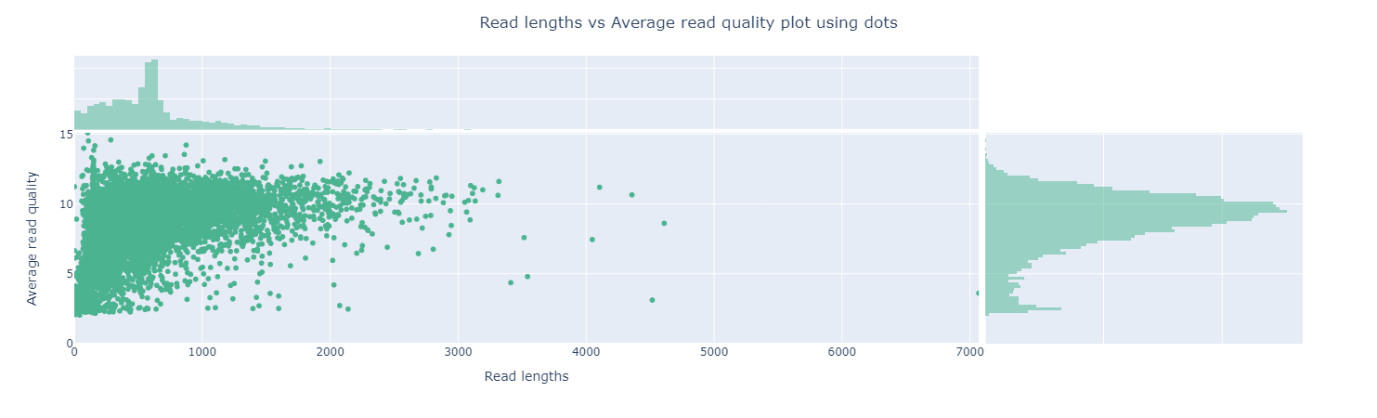

(a)

(b)

**Fig S1**: Read length vs average read quality dot plot (a) PFC (b) PVC

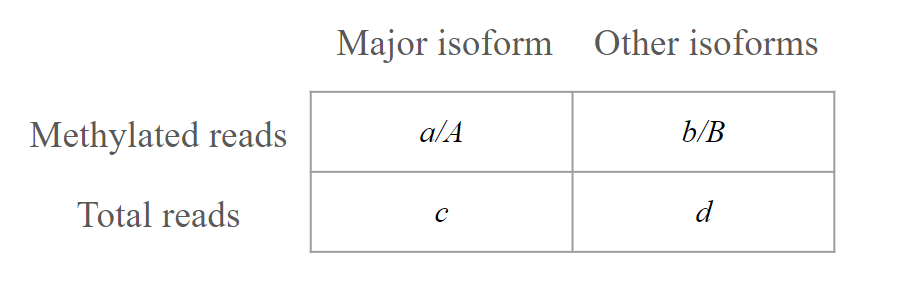

**Fig S2**: 2X2 contingency table to perform Fisher's test to identify genes displaying differential splicing of exons.

**Supplementary Tables**

| **Features** | **Sample Details** | |
| --- | --- | --- |
| Sample Code | PFC | PVC |
| Parasite Density | 100000/µL | 40000/µL |
| Microscopic Investigation | Ring 100% | Troph 60% + Schizont 40% |
| Hb | 5.2 | 6.8 |
| Serum bilirubin | 11 | 3.6 |
| Starting Qubit Concentration | 1 ug total RNA | 1.2 ug total RNA |
| Clinical Presentation | J (ser. bil. - 11), A (Hb. – 5.2) | J (ser. bil. - 3.6), A (Hb. - 6.8) |

Table S1: Clinical characteristics of patient samples used in the study. J: Jaundice; ser.bil: serum bilirubin(mg/dL); A: Anemia; Hb:Hemoglobin(g/dL)

| **Feature** | **PFC** | **PVC** |
| --- | --- | --- |
| Number of reads | 207,945.0 | 672,843.0 |
| Total bases | 290,301,217.0 | 124,014,603.0 |
| Mean read length | 596.4 | 431.5 |
| Mean read quality | 7.4 | 7.6 |
| Median read length | 553.0 | 467.0 |
| Median read quality | 9.1 | 8.9 |
| Read length N50 | 676.0 | 551.0 |
| mean depth (genome) | 3.6x | 0.9x |
| % of reads above quality cut-offs (Q>7) | 81.5% | 83.3% |
| Mean Quality Score | 15.7 | 15.8 |
| Number of processed reads | 169464 | 560535 |
| %GC | 23.44% | 44.8% |
| Number of full-length (FL) transcripts reconstructed | 73282 | 40341 |
| Number of full-length (FL) transcripts reconstructed (without ISM) | 66,102 | 36,926 |
| Number of unique isoforms | 3258 | 3130 |

Table S2: Sequencing and Transcriptome statistics of PFC & PVC sequencing experiment.
